# Graph-Dynamo: Learning stochastic cellular state transition dynamics from single cell data

**DOI:** 10.1101/2023.09.24.559170

**Authors:** Yan Zhang, Xiaojie Qiu, Ke Ni, Jonathan Weissman, Ivet Bahar, Jianhua Xing

## Abstract

Modeling cellular processes in the framework of dynamical systems theories is a focused area in systems and mathematical biology, but a bottleneck to extend the efforts to genome-wide modeling is lack of quantitative data to constrain model parameters. With advances of single cell techniques, learning dynamical information from high throughput snapshot single cell data emerges as an exciting direction in single cell studies. Our previously developed dynamo framework reconstructs generally nonlinear genome-wide gene regulation relations from single cell expression state and either splicing- or metabolic labeling-based RNA velocity data. In this work, we first developed a graph-based machine learning procedure that imposes a mathematical constraint that the RNA velocity vectors lie in the tangent space of the low-dimensional manifold formed by the single cell expression data. Unlike a popular cosine correlation kernel used in literature, this tangent space projection (TSP) preserves the magnitude information of a vector when one transforms between different representations of the data manifold. Next, we formulated a data-driven graph Fokker-Planck (FPE) equation formalism that models the full cellular state transition dynamics as a convection-diffusion process on a data-formed graph network. The formalism is invariant under representation transformation and preserves the topological and dynamical properties of the system dynamics. Numerical tests on synthetic data and experimental scRNA-seq data demonstrate that the graph TSP/FPE formalism built from snapshot single cell data can recapitulate system dynamics.

**Significance Statement:** A cell is a dynamical system, which responds to extracellular and intracellular cues and changes its internal state. In practice the internal state of a cell is often characterized by its gene expression profile such as its transcriptome measured through destructive single cell techniques. Just like one can use Newton’s equations to describe motions of the celestial bodies in the solar system, the state change of a cell in principle can also be described by a set of mathematical equations. Determining the form and associated parameters of such equations, however, is challenging. This work presented a general framework of reconstructing dynamical equations from snapshot single cell data.

## Introduction

Cells undergo transitions between different states, each characterized by distinctive expression profiles of their transcriptome, proteome, and epigenome, among other factors. The study of cellular dynamics within the framework of dynamical systems theory has been a focused topic in mathematical and systems biology. However, these studies have often been restricted to modeling a limited number of selected molecular species. As these molecular species are components of a large interconnected cellular network, an important open question is how the dynamics of the subsystem under study couple to the remaining part of the network.

During major cell fate or type transitions, cells undergo significant changes in their global expression profiles. Traditional bottom-up systems biology approaches face challenges in investigating how different cellular programs function coordinately during such transitions. To address these challenges, it is crucial to explore how various cellular processes and molecular components interact to orchestrate the overall behavior of the cell.

Recent advances in single cell techniques such as scRNA-seq and scATAC-seq provide genome-wide albeit snapshot information on cell states. Extraction of dynamical information from single cell snapshots recently emerged as an active research area (1-4). Notably, La Manno et al. showed that one can infer gene-specific instant changes in the mRNA levels called RNA velocities from scRNA-seq data (5). This seminal study has led to numerous efforts to further improve the accuracy of the velocity estimation using either the original splicing-based model or more direct measurements of mRNA turnover dynamics with metabolic labeling approaches. Using these splicing- or labeling-based discrete single cell expression and RNA velocities as input, Qiu et al. developed a framework for reconstructing the genome-wide and generally nonlinear dynamical models of gene regulatory networks governing cellular processes (**Fig. 1A**) (6). For example, they reconstructed the differential sigmoidal- and exponential-shaped dose-response relations between two antagonistic master regulators GATA1 and SPI1 during hematopoiesis that agree with previous experimental measurements.

**Figure 1.**
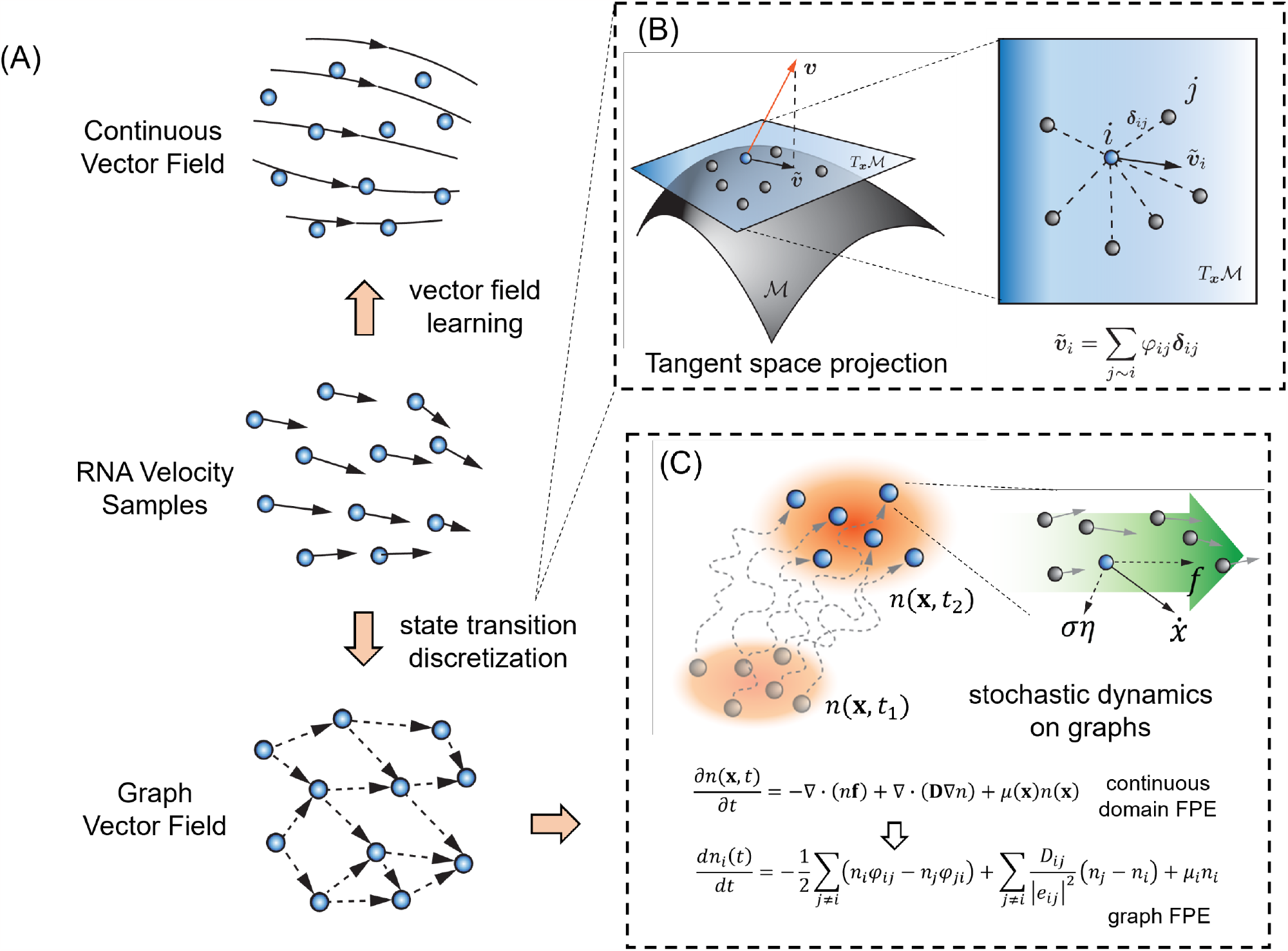
Discretization of RNA velocity vectors in state transition graphs. **(A)** Two ways of modeling dynamics from single cell RNA velocity data. *Top*: Reconstruction of a continuous vector field using vector field learning algorithms, as presented in (6). *Bottom*: Discretization of RNA velocity vectors into state transition flows on the neighborhood graph with cells as vertices. **(B)** *Left*: The tangent space ***T***_**x**_ℳ (*blue* plane) of a data manifold ℳ (curved surface) at the point **x**. The *red* arrow represents the unprojected RNA velocity vector. The *gray* spheres represent neighboring cells of the *blue* cell. *Right:* The neighbors of **x** form a neighborhood graph that can be used to approximate the tangent space locally. 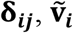distance vector and projected velocity vector in ℝ^***d***^. **φ_*ij*_**: cell transition flows and also components of the graph vector field. **(C)** Stochastic modeling of cell dynamics. *Left*: The propagation of cell population, described by the evolution of cell density distribution ***n***(**x, t**), over time. *Right*: decomposition of RNA velocity vector into deterministic (**f**) and stochastic components (**ση**). The *green* arrow in the background indicates the collective flow around the *blue* cell.

While these studies have demonstrated the feasibility of reconstructing cellular dynamics from snapshots, further developments are needed for improving the accuracy of the predictions. Specifically, for dynamo there are several areas that can be further developed:

1. Imposing mathematical constraint of a dynamical process on the RNA velocity estimation with physics-informed machine learning (7). Dynamo assumes that the measured single cell expression profiles and inferred RNA velocities collectively reflect a dynamical process and are connected through a set of dynamical equations. Existing methods of estimating single cell RNA velocities have not taken into account such constraint.
2. Accounting for the complete stochasticity of the cellular dynamics in a cell state space. Cellular dynamics is stochastic and can be genetically modeled as, e.g., a convection-diffusion process (1, 8). Dynamo and a subsequently developed DeepVelo approach (9) reconstruct the convection part only.
3. Efficiently organizing and analyzing data embedded in a high-dimensional space. Conventional numerical methods study dynamical systems using lattice grids that represent the ambient space, but the number of grids increases exponentially with the dimensionality of the space, a phenomenon known as the curse of dimensionality.
4. Transforming the dynamical equations between different representations and projecting them onto a relevant subspace that preserves topological and dynamical features of the system dynamics. Numerous algorithms have been developed in data science to transform between different representations and perform dimension reduction. These algorithms are designed to deal with static data without considering the underlying dynamical processes, or the fundamental requirement that physical laws are independent of the underlying coordinate system and the physical equations should be covariant under coordinate transformation. One example is the invariance of time, mass, and acceleration of a classical particle under a Galilean transformation between two reference frames moving relative to each other in the nonrelativistic regime. In the context of single cell dynamics, for representing an RNA velocity vector in different representations, a heuristic and empirical cosine kernel approach is widely used by constructing an RNA velocity-related Markov transition model between neighboring cells (5). This approach is intuitively appealing but lacks mathematical foundation, and its limitations have been discussed in the literature (10). A mathematically rigorous approach is needed.

In this study, we tackle the above challenges through mapping the cellular dynamics onto a discrete graph representation. We first propose a simplistic approach to describe the cell state transitions using a graph-theoretical representation of RNA velocities with biophysical and dynamical systems underpinnings (**Fig. 1A**). Then, we model the stochastic dynamics based on the state transition graph as a convection-diffusion system, using a graph discretization of the continuous Fokker-Planck equation (FPE) together with a tangent space projection (TSP) procedure for imposing the dynamical constraint (**Fig. 1B** and **C**). We benchmark the proposed graph TSP/FPE framework with both simulated experimental single cell data.

## Results

### Projecting velocity vectors to the tangent space from single cell RNA velocities using displacement vectors of a neighbor graph

Assume that one can identify a set of dynamical variables to fully specify a cell state. With single cell transcriptomic data composed of *d* genes, we further assume that the mRNA count of each gene (after proper normalization if needed), organized as the elements of a *d*-dimensional vector **x**, is proportional to its cellular concentration and can serve as the dynamical variables forming an ambient multi-dimensional gene expression space, ℝ^*d*^. After transient relaxation of dynamical degrees of freedom faster than the temporal resolution under study, the temporal evolution of a dynamical system can be typically described by changes along an *m*-dimensional manifold ℳ embedded in ℝ^*d*^ with *m* ≪ *d* (11). Indeed, it is widely established in practice that high dimensional data, such as those inferred from scRNA-seq experiments, reside in a curved space, known as the *data manifold*, whose dimension is much lower than that of the ambient space where it is embedded. The framework can be easily generated to a set of coupled stratified manifolds corresponding to, for example, different epigenetic states.

With cells as dynamical systems, if the dynamics is continuous, the evolution of **x** along the manifold ℳ, can be described by defining a velocity vector related to an infinitesimal propagation operator that maps the cell state **x** at time t to **x**’ at a later time, both on the manifold ℳ, 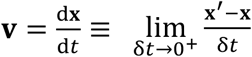. The velocity vectors for all genes form a vector field on ℳ, and at any given state **x** the velocity vector **v** lies in the space *T*ℳ (**Fig. 1B**), which is tangent to the manifold ℳ. We denote the tangent plane at a point **x** on ℳ as *T**_x_***ℳ.

With sufficient sampling size, we assume that the manifold ℳ can be well approximated by the data 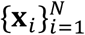, where *N* is the number of cells. On the other hand, the RNA velocity vector of a cell **v** is derived from either splicing or metabolic labeling data generally with less accuracy (5, 6). Consequently, in practice the RNA velocity vectors estimated from experimental datasets do not always reside in the tangent space of the observed data manifold. They may also have a component normal to the tangent plane which may originate from various sources such as inaccurate measurements of unspliced/spliced or labeled/unlabeled RNA species, intrinsic errors arising from RNA velocity estimation methods (6, 10, 12), and/or incomplete representation of cell states in the transcriptomic space.

We hypothesize that one can correct the measured RNA velocities **v** using the more reliable data manifold ℳ and the mathematical constraint that **v** should lie in the tangent space *T*ℳ. First, we notice that the infinitesimal neighborhood of a given point **x**_*i*_ corresponding to a sampled cell *i* can be approximately represented by a Euclidean space defined by **x**_*i*_ and the states of neighboring cells. With sufficient sampling of the cells *j* neighboring to cell *i* in the state space, the distance vectors between cell *i* and its neighboring cells, **δ**_*ij*_, form a set of complete albeit possibly redundant and nonorthogonal/non-normalized basis vectors of the Euclidean space. Then the projection of the measured velocity vector **v**(**x**_*i*_) of cell *i* onto the tangent surface *T*_**x**_ℳ, denoted as **v**_‖_(**x**_*i*_), can be expressed as a linear combination of **δ**_*ij*_,

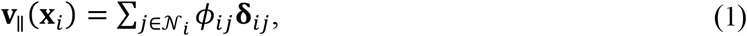

where 𝒩_*i*_ is the neighborhood of cell *i*, defined by its *k* nearest neighbors in the expression state space. Here *ϕ*_*ij*_ scales with the cosine of the angle between **v**(**x**_*i*_) and **δ**_*ij*_. Direct application of Eqn. 1 to determine the coefficients is numerically unstable (see ST Section A for detailed discussions). Instead we determined 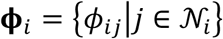 by minimizing the following loss function,

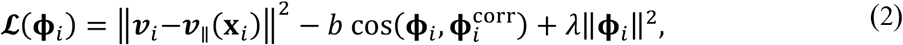

where ||.|| refers vector modulus. Here 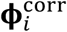 is a heuristic “cosine kernel” widely used in the RNA velocity community for embedding the RNA velocity vectors in a reduced space (5, 13), *b, λ* are two hyperparameters determining the emphases on retaining the direction of 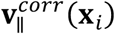 and regularization, respectively. The L2-regularization is used to bound ‖**ϕ**_*i*_‖.

In the above formulation, we re-casted the cosine kernel in the context of the tangent space projection with a form similar to that of Eqn. 1:

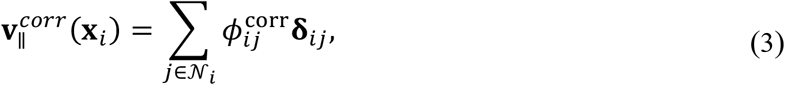

where 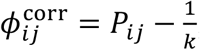,and 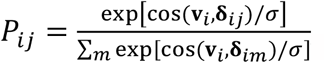 with cos(·,·) denoting the cosine similarity between two input vectors, *σ* an arbitrary bandwidth parameter, and *k* the number of neighbors for each cell. Here *P*_*ij*_ gives a heuristic transition probability from cell *i* to *j*. The term (−1/*k*), called the “density correction”, is designed to correct the potential sampling bias where the embedded velocity vectors tend to point towards the direction of regions with high cell density. Notice that the correlation kernel loses information about the magnitude of the velocity vectors **v**_*i*_ (i.e., the *speed*), due to the normalization in the correlation functions. Therefore, the correlation kernel is empirical without rigorous mathematical foundation, and qualitatively guided by the physical intuition that a cell has a high tendency to move along the direction of its velocity vector. Here, we used such physical intuition to help on determining **ϕ**_*i*_ uniquely.

#### Representing scalar and vector fields on a discrete and directed graph preserves invariance in topological structures of a manifold

In the above section we considered how to represent a vector at a single point of a manifold using its neighborhood graph. In this section, we further formulate a discrete representation of a scalar or a vector field defined on a continuous manifold.

First, we approximate a continuous manifold by a discrete network formed by the sampled data points in the ℝ^*d*^ gene expression space. Through Eq. 1, a cell state *i* and those in its neighborhood separated by displacement vectors **δ**_*ij*_ form a weighted local network. Each **δ**_*ij*_ is defined by its magnitude, sense (i.e., from **x**_*i*_ to **x**_*j*_), and orientation (as characterized by the angle between **δ**_*ij*_ and a reference vector). All the data points and their neighborhood displacement vectors form a network approximating the continuous manifold ℳ.

Next, we map the network homeomorphically to an abstract graph *Γ*(*Ω, E*), where *Ω* and *E* are the sets of vertices and edges with one-to-one correspondence to those of the network in the ℝ^*d*^ space. Therefore, the network of (**x**_*i*,_**δ**_*ij*_) is a finite neighborhood approximation of the tangent space of a continuous manifold ℳ at site **x**_*i*_, and the corresponding portion of the graph *Γ* provides an abstraction in the graph representation (**Fig. 1B**).

To understand the advantage of working with the graph space, consider embedding the data manifold in an ambient space different from the original ℝ^*d*^ space. The two representations are connected through a homeomorphic transformation between (**x**_*i*,_**δ**_*ij*_) and 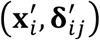. Geometric features of both the continuous manifold and the network as its discrete representation, such as curvatures of the manifolds and angles between two displacement vectors, are representation-dependent. On the other hand, since the same set of datapoints are used to approximate the manifold in different representations, both the connectivity of vertices and the sense (i.e., identities of the starting and ending points) of a displacement vector are invariant to the representation transformation. A graph representation preserves the topological invariance, while neglecting the representation-dependent geometric characters. This topological abstraction is achieved by mapping each displacement vector **δ**_*ij*_ to a directed edge *e*_*ij*_ on a graph, with a measure of magnitude and sense but not orientation. Consequently, a class of topologically invariant manifold structures map to the same graph *Γ*.

With the graph *Γ*, one can define scalar and vector functions on *Ω* and *E*, respectively (13, 14). While scalar functions are defined on vertices, vector functions are defined on the edges of the graph. For *i, j* ∈ *Ω* and *e*_*ij*_ ∈ *E*, the edge connecting the two vertices, a single graph vector **ϕ** at edge *i* can be defined as, 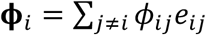, which corresponds to Eqn. 1 in the ℝ^*d*^ space. Then a vector field forms a distributed directional flow on the graph through a collection of graph vectors. Consequently, *ϕ*_*ij*_ and *ϕ*_*ij*_ describe the same probability flow along the edge (*e*_*ij*_ = −*e*_*ij*_), with *ϕ*_*ij*_ = −*ϕ*_*ij*_. In practice, *ϕ*_*ij*_ and (−*ϕ*_*ij*_) estimated from two neighboring data points (e.g., from Eqn. 2) may not be the same, then one can take their mean as an estimation. Therefore, a vector field specifies non-negative and unidirectional flows on the edges 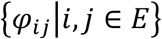, which can be estimated from datapoints through,

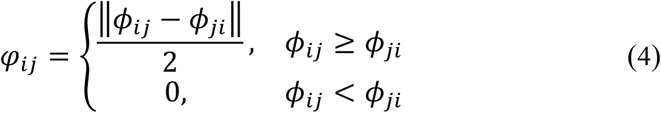

### Stochastic dynamical equations of cell dynamics on a graph are covariant under homeomorphic transformation

Consider that the dynamics of a cellular process can be described by the following generic stochastic differential equations (SDE) on the manifold ℳ:

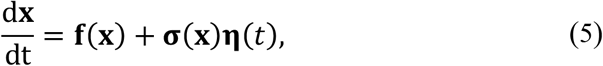

with a drift/convection term **f** and a diffusion term **ση**, where ⟨*η*_*i*_(*t*)⟩ = 0 and ⟨*η*_*i*_(*t*)*η*_*j*_(*t*′)⟩ = δ_*ij*_δ(*t* − *t*′), where the former δ_*ij*_ is the Kronecker delta, and the latter δ(*t* −*t*′) the Dirac’s delta function. The average is over all realizations of the noises. Eqn. 5 describes the state evolution of a tagged cell. A cell in general also undergoes proliferation or death, quantified by a rate constant *μ*(*x*), which requires a kinetic equation separate from Eqn. 5.

Averaging of Eqn. 5 over the noises at a fixed **x** gives 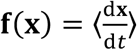. Notice that the reported single cell RNA velocities are raw RNA velocities estimated from the splicing or labeling data then averaged over neighboring cells, which can be taken as a replacement for the noise average in practice. Then with Eqn. 1, at a sample point **x**_*i*_,

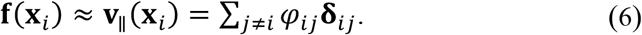

From the population perspective, we can define a cell density function *n*(**x**, *t*) in the expression space. The temporal evolution of the population density corresponding to Eqn. 5 is described by a set of Fokker-Planck equation (FPE) (15, 16),

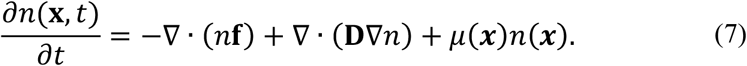

Here we assume *μ*(***x***) as a localized function (see (1) for more general treatment), and 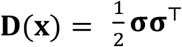 is the diffusion tensor. Similar mathematical framework has been used to model other processes such as protein motor dynamics (17, 18).

To solve Eqn. 7 numerically, one can divide the cell state space into subregions, e.g., using square lattices, and denote *n*_*i*_ the cell number within subregion *i*. Then Eqn. 7 can be converted into a discrete kinetic model of transitions between subregions (17, 18). A challenge is that the needed number of lattice cells grows exponentially with the dimensionality. A solution we propose here is to work on the discrete graph corresponding to the generally low-dimensional manifold ℳ directly. Analogous to the familiar continuous calculus, one can define differential operators on a graph(19) (see **ST Section B-D** for details). Then it is straightforward to convert Eqn. 7 to its counterpart on a graph,

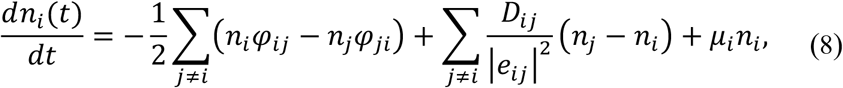

where the diffusion tensor **D** becomes a function of graph edges. One can further write the above equation in a vectorized form:

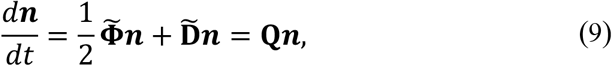

where,

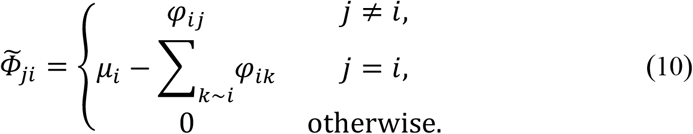

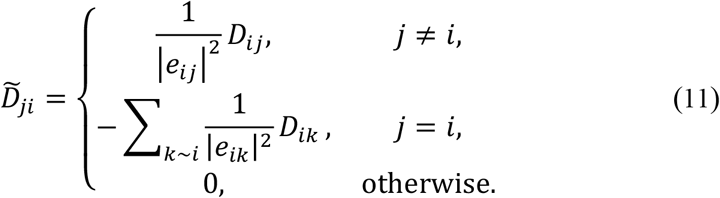

Physically, each vertex represents a subregion with irregular shape generally, and *φ*_*ij*_ gives the net convection flux density through the boundary surface between two neighboring subregions. Once the subregions are divided, while the shape of each subregion changes along with homeomorphic representation transformation, quantities such as *n*_*i*_, *φ*_*ij*_, *μ*_*i*_, and *t* need to be invariant, following the principle of covariance of physical laws. While there is freedom on choosing *e*_*ij*_, one can rescale the diffusion tensor element *D*_*ij*_ so the reduced diffusion matrix term 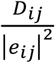 with a dimension of 1/time remains invariant, and the overall dynamical equation is invariant under representation transformation. Therefore, the matrix 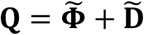 is the transition rate matrix for a continuous time Markov chain (CTMC) defined on the graph, and the transition rate *Q*_*ij*_ is defined as the probability of the (convectional and diffusive) transition from vertex *i* to *j* per unit time (see also **ST Section E**).

Equations 8-11 are the main theoretical results for reconstructing dynamical equations on a graph from single cell genomics data. In the case having RNA velocities available, Eqn. 6 provides one practical approach of determining *φ*_*ij*_. With additional information, such as scRNA-seq data at different time points and lineage tracing data, one can also constrain other parameters 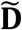 and μ following procedures similar to what used in other studies (2).

### Tangent space projection preserves magnitudes of the velocity vector field

To demonstrate the effectiveness of using RNA velocity-based graph vector fields to model cell state transitions, we first considered two systems: (1) a two-gene toggle switch, and (2) simulated neurogenesis. Both models were adapted from ordinary differential equation (ODE) models used in previous studies (20) and were simulated using the Gillespie algorithm (see **ST Section F** and **G** for details).

The two-gene toggle switch model consists of two mutually inhibited, self-activating genes, modeled with inhibition and activation Hill functions, respectively (**Fig. 2A**). Under the chosen set of kinetic parameters, there are in total three fixed points, including a saddle point in the middle, and two attractors symmetrically distributed on its two sides. The cells are initialized along the separatrix with equal distances from the two attractors and simulated to migrate to the attractors using the Gillespie algorithm for a sufficiently long period of time. The neurogenesis model, on the other hand, consists of 12 genes forming a network that contains two consecutive toggle switch motifs, e.g., the mutual inhibition between *Mash1* and *Hes5*, activated by *Pax6*; and between *Scl* and *Olig2*, activated by *Hes5* (**Fig. 2B**) (20). The network has three attractors corresponding to three stable cell types: neuron, astrocyte, and oligodendrocyte. All genes initially have zero expression levels, except for *Pax6*, whose expression level is determined based on the bifurcation analysis from Qiu et al. (20). For both models, 5,000 cells were sampled from ten simulated trajectories, and the reactions rates from the corresponding ODE models were used as the “ground truth” RNA velocity vectors.

**Figure 2.**
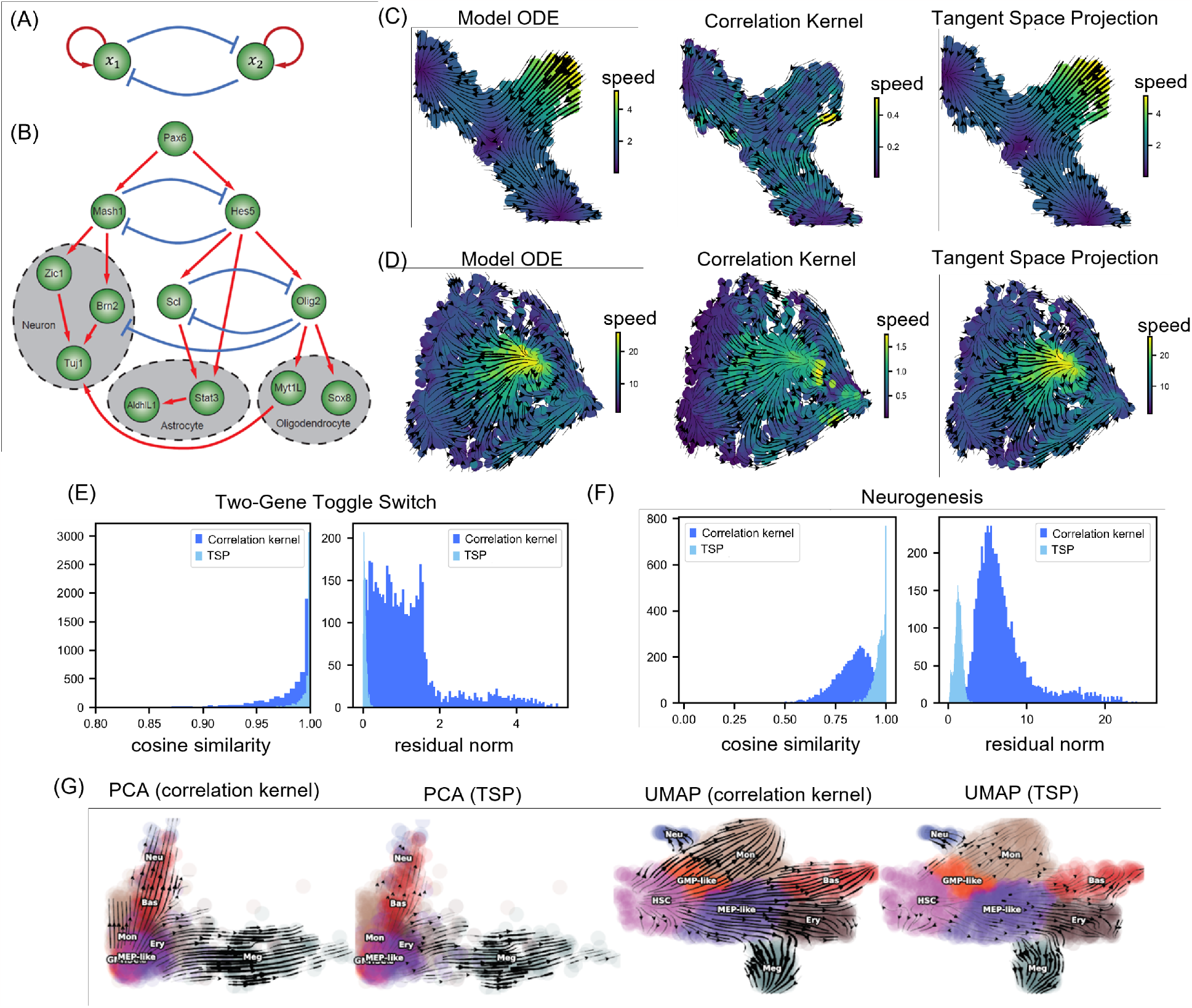
Two-dimensional velocity projection of RNA velocity vector fields reconstructed from simulated and scRNA-seq datasets. (A) The two-gene toggle switch model (see **ST Section F** for the ODEs). (B) The Neurogenesis model with 12 genes and three mature cell lineages (see **ST Section G** for the ODEs). (C) Comparison between the velocity vectors from the ODEs of the two-gene toggle switch model (*left*, ground truth) and velocity vectors reconstructed from simulated data then projected to the two-gene expression space by the cosine correlation kernel (*middle*) or co-optimization generated graph vector field (*right*). (D) Similar as panel C but for the neurogenesis model, with reconstructed velocity vectors projected to the leading two-dimensional PCA space. The ground truth vectors are the first two components of the 12-dimensional velocity vectors in the PCA space, which are obtained through rotating the velocity vectors in the original gene space specified by the ODEs. (E) Histograms of cosine similarity (*left*) and L2-norm of residuals (*right*) between model ODE velocity vectors (ground truth) and velocity vectors projected using the correlation kernel and co-optimization generated graph vector field for the two-gene toggle switch model. (F) Similar as panel E but for the neurogenesis model. (G) Two-dimensional projection of RNA velocity vector fields of the human hematopoiesis dataset in the leading PCA and UMAP representations, respectively.

For each model, we used two methods to project the RNA velocity vectors onto the same two-dimensional space (**Fig. 2C & D**). As expected, the correlation kernel projected velocity vectors reasonably capture the direction but not the magnitudes of the true RNA velocity vectors for both models. In comparison, the velocity using the TSP projection on the PCA space preserves both the direction and the magnitude of the original RNA velocities. To quantify the different performance of these two projection methods, we also calculated the cosine correlation as well as the L2 norm of the residue between a projected vector and a ground truth vector (**Fig. 2E and F**). For TSP, both the cosine correlation and the residue norm have a narrow distribution peaked around one and zero, respectively, indicating close agreement between the reconstructed and the true velocity vectors. The results of the cosine kernel vectors, on the other hand, have much broader distributions with peaks shifted away from one and zero, respectively.

Next, we compared the two projection methods using previously acquired hemopoietic stem cell differentiation and murine pancreatic endogenesis datasets(6, 21). In these real data cases the ground truth is not available. In both PCA and UMAP representations, the streamline plots of the projected velocity from both methods accurately revealed that hematopoietic progenitor cells first bifurcate into either the GMP or MEP lineages and then forms ramifications leading to five terminal cell fates (**Fig. 2G**), and ductal cells differentiate into alpha, beta, gamma, and epsilon cells through a number of intermediate cell types during pancreatic endogenesis (**Fig. S1A** and **B**). A parameter scan analysis (**Fig. S1C**) shows that for the two parameters in Eqn. 2, *b* controls local smoothness and *a* controls how well the velocity magnitude is conserved. For both datasets, there are noticeably quantitative differences between results from the cosine kernel and the TSP projection methods. These results agree with our conclusions with the simulated model systems that the TSP but not the cosine kernel projection methods can preserve both the directions and magnitudes of a vector field.

### Graph FPE accurately captures the kinetics of cell differentiation

It should be noted that the transition matrix constructed using the correlation kernel (Eqn. 3) models the cell dynamics qualitatively, in the sense that its source and sink nodes correspond to the repulsors and attractors of the modeled vector field **f**. The stationary distribution of the resulted discrete time Markov chain (DTMC), and properties of which, e.g., fate probabilities, generally reflects cell fate decisions given by the directional information of RNA velocity vectors. However, because of the removal of the speed and heuristic construction of the transition probabilities, the kinetics of the DTMC is not comparable to that of the FPE described here. Therefore, it is not feasible to infer any time-related properties of the underlying cell dynamics from the correlation kernel constructed DTMC. That is, since the cosine kernel does not preserve the magnitudes of velocity vectors, the resultant transition matrix may not recapitulate even the qualitative information of the sequence of competing events that take place in parallel in a cellular process.

To examine how well the TSP method and the associated graph FPE can capture the kinetics of cell state transitions, we performed CTMC simulations by analytically solving Eqn. 9 to propagate states of cells sampled from an initial distribution using the transition matrix for the two-gene toggle switch model (**Fig. 3A**). The graph FPE correctly describes the temporal evolution of cells moving away from the initial positions to the central saddle point, and then bifurcate into the two attractors.

**Figure 3.**
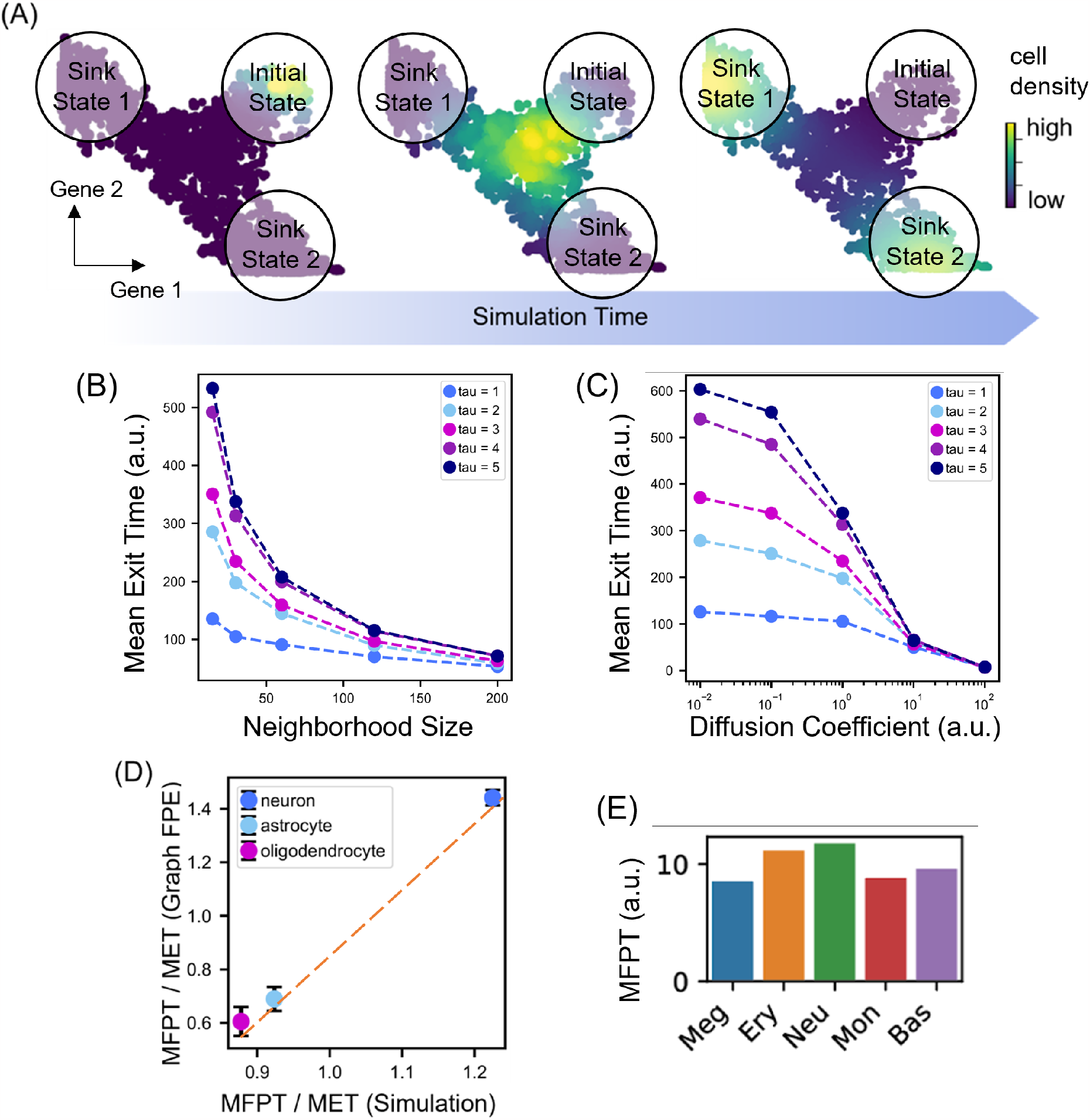
Graph FPE quantitatively recapitulates the stochastic kinetics of of a cellular process. **(A)** Propagation of cell density distribution ***n***(**x, *t***) modeled using graph FPE for the simulated two-gene toggle switch model. The color represents the value of the density at each cell state, with brighter color indicating higher probability. The sum of probability at all cells is one. **(B, C)** Mean exit time calculated from graph FPE for the two-gene toggle switch model of different kinetics (controled by the time scale paramter **τ**) affected by the (B) neighborhood size, when the diffusion coefficient is fixed as one; and the (C) diffusion coefficient, when the neighborhood size is fixed as 30. **(D)** Graph FPE captures the distinctive kinetics of cell state transitions from naïve cells to three mature cell types in the simulated neurogenesis model. The *x*-axis is the ratio of MFPT to MET calculated from the simulated trajectories using Eqn. S37 and S38. The *y*-axis is the ratio of MFPT to MET calculated from the graph TSP/FPE using Eqn. S39 and Eqn. S40. (E) MFPT of various differentiated cell states calculated from the graph FPE model of the human hematopoiesis dataset (6).

The graph FPE equation in Eqn. 8 indicates that besides the convection vector field and the birth-death rate, the dynamics of a process is also influenced by the graph neighborhood size and the diffusion constant. To examine how these parameters affect the dynamics of a graph FPE model, we performed CTMC simulation on FPE models reconstructed from simulated data of a panel of two-gene toggle switch models, which do not include the birth-death process. These toggle switch models assume parameters that lead to the same stationary distribution and differ only by a timescale scaling parameter τ (see **ST Section F**). A smaller τ gives rise to faster kinetics, i.e., the trajectories reach the two attractors within a small simulation time interval, and with larger RNA velocity vectors. To quantify the time needed for simulated cells to migrate from the initial position to the two attractors, we calculated the *mean exit time* (MET) from the simulated trajectories and the constructed CTMC. MET is defined as the average time for the cell to reach *any* target state from the initial position for the first time. We designed these models since a graph FPE model constructed from the cosine kernel, i.e., replacing 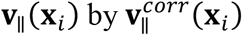 in Equation 6, would get the same MET for the systems with different τ and thus cannot distinguish fast and slow dynamics. The results in **Fig. 3 B** and **C** show that the graph FPE models robustly recapitulate an increasing MET with a slower convection dynamics (i.e., a larger value of τ). With a fixed diffusion constant, the calculated MET of a model system with a smaller τ numerically converges faster with the neighborhood size as expected (**Fig. 3B**). This is because a smaller neighborhood leads to less accurate description of diffusion, but the resultant overall numerical error is less when convection weights more on diffusion to contribute to the system dynamics. When the diffusion constant increases, diffusion eventually dominates, and the MET of different models converges to a same value (**Fig. 3C**). These results indicate that while the neighborhood size and the diffusion constant affect the precise value of a dynamical quantity calculated from a graph FPE model and should be further constrained (e.g., by temporal evolution of the single cell state distributions), the results are robust on providing qualitative and even semi-quantitative information on the relative dynamics between branched cellular events insensitive to the parameter choice.

To further test whether the graph TSP/FPE model can be used to determine the temporal order of cell commitment to different cell types during a differentiation process, we applied it to the simulation data generated from the neurogenesis model. To characterize the expected time for cells to reach each stable cell type, we calculated both the MET, and the *mean first passage time* (MFPT) for the three cell types. While an MET is defined as the expected time to reach any of the final states (i.e., the three stable cell types), an MFPT is the expected time to reach a specific target state before reaching any other sinks (see **ST Section H & I** for details). Both the Gillespie simulations of the neurogenesis model and CTMC simulations of the FPE models show that the oligodendrocyte appears the earliest, followed by the astrocyte, and it takes the longest time for cells to become mature neurons, and the MFPTs calculated from the graph FPE for the three cell types agree with those calculated from the Gillespie simulations of the original model **(Fig. 3D)**. The sequence of appearance of the three cell types is robust when we vary the diffusion constant and the neighborhood size. Similarly, MFPT analyses on the graph FPE model reconstructed from the human haemopoietic stem cell differentiation dataset predict that Meg cells appear the first (**Fig. 3E**), agreeing with experimental observations (22). These results further support that the graph TSP/FPE model is capable of predicting temporal sequence of parallel cellular events.

#### A graph Vector field can be approximated as the gradient of a scalar field under some conditions

The Helmholtz-Hodge decomposition theorem states that any vector field can be decomposed into three parts (**Fig. 4A**): the gradient of a scalar potential, the curl of a vector potential, and a harmonic function (23, 24). On reconstructing cellular dynamics from single cell snapshot data, for numerical simplicity one often uses a scalar potential to approximate the governing vector field (2, 25). In this case the velocity components are no longer independent but can be calculated by as the gradient of the scalar potential. This property is especially convenient when the spliced or labeling RNA velocity are not available or reliably estimated for a system or for a subset of genes, but can be derived from a scalar potential.

**Figure 4.**
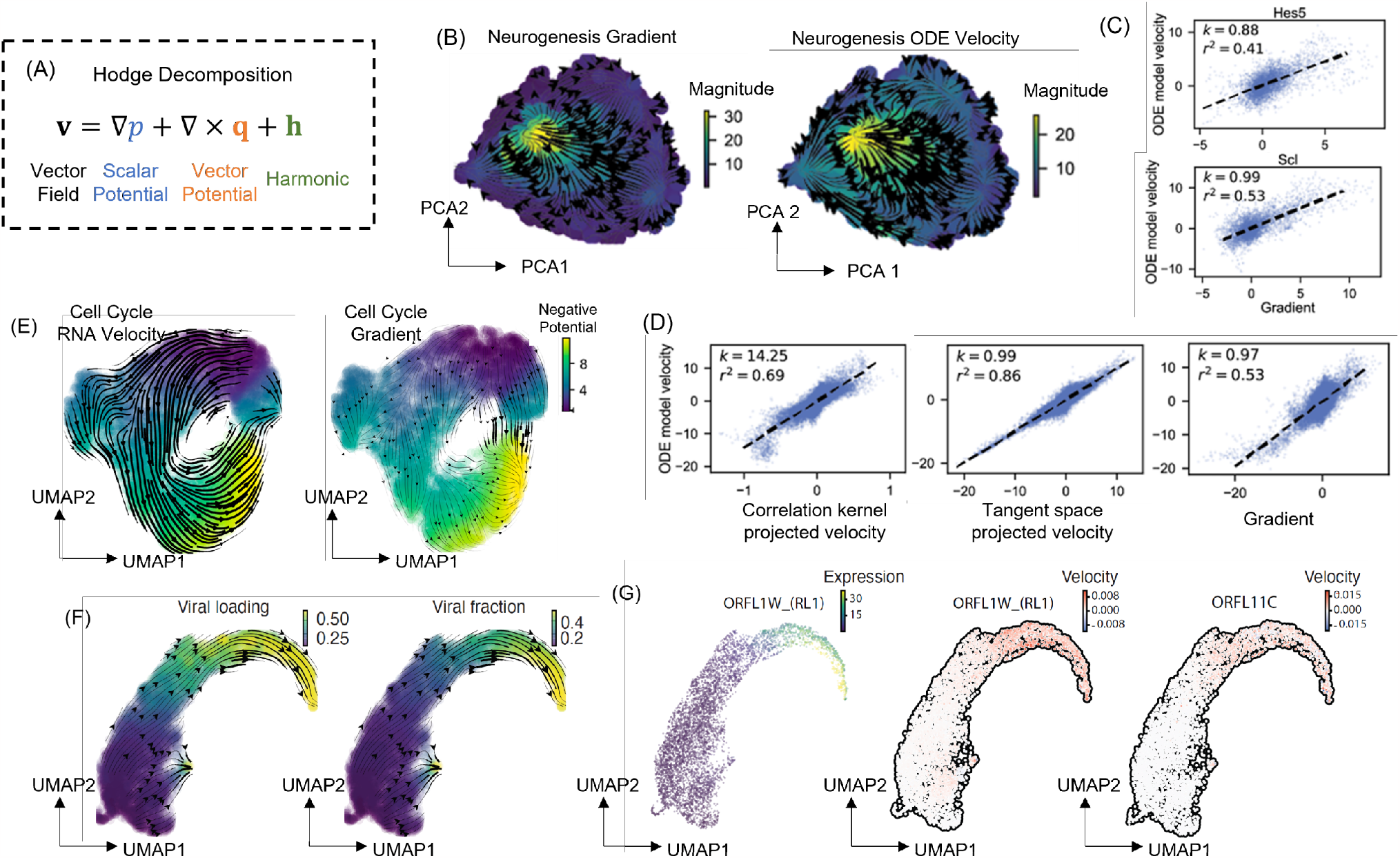
A vector field can be approximated as the gradient of a scalar potential under some conditions. (**A**) Schematics of the Helmholtz-Hodge decomposition of vector fields. Any vector field can be decomposed into the gradient of a scalar potential (*blue*), the curl of a vector potential (*orange*), and a harmonic vector field (*green*). (B) Streamline plots of the gradient based RNA velocity and the true RNA velocity projected in the PCA space. (**C**) High concordance of the gradient based RNA velocities and the true RNA velocities across cells of two key regulators, *Hes5* and *Scl* in the neurogenesis simulation system. (**D**) Correlations of correlation kernel projected velocities (*left*), TSP kernel projected velocities (*middle*), and gradient based velocities (*right*) with respect to the ground truth velocities from the neurogenesis ODE model. (**E**) RNA velocity (*left*) and gradient based velocity streamlines (*right*) of human A549 cell cycle process captured by scNT-seq (29). (**F**) The phase plot of the predicted RNA velocity with the graph gradient operator using viral loading (*left*) and viral fraction (*right*) as potentials. (**G**) The scatter plot of the gene expression (*left*) and predicted RNA velocity across cells with the graph gradient operator of viral genes ORFLW_(RL1) (*middle*) and ORFL11C (*right*).

We tested the scalar potential approximation through applying a discrete Hodge decomposition algorithm and the graph gradient operator on graph velocity vector field reconstructed from both simulated and experimental data. For the simulated neurogenesis system, a direct visual inspection already shows that there is high consistency between the streamlines of the gradient operator-based velocity and that of the ground truth velocity (**Fig. 4B**). We observed similar consistency on the velocities at the gene level (**Fig. 4C**) or projected velocity in the PCA space (**Fig. 4D**). The RNA velocity projected to the PCA space using the gradient based approach has a narrow distribution of the cosine similarity to the original data, with the peak close to one. The results indicate that in this case the velocities estimated from the gradient approximation are comparable to that calculated from the full vector field using the TSP kernel, and the approximation even outperforms the correlation kernel-based projection method which does not preserve the magnitude of the vector field as discussed earlier (**Fig. 4E**). Next, we analyzed a cytomegalovirus (CMV) viral infection dataset (26) to learn the viral transcriptomic kinetics (**Fig. 4F**) that does not involve RNA splicing. Interestingly, RNA velocities derived as the gradient of the viral loading and fraction reveals the expected temporal progression of viral accumulation (**Fig. 4G**).

Given the encouraging results of approximating a vector field as the gradient of a scalar potential, one should keep in mind that the approximation breaks down when the non-gradient terms become non-negligible. One such case is when cell cycle is present. Indeed, analyses on a metabolic labeling dataset of proliferative A549 cells treated with dexamethasone reveal noticeable differences between the full vector field streamlines and those from the gradient approximation (**Fig. 4E**). This result emphasizes caution on applying the gradient approximation.

## Discussion

In this work, we developed a mathematically rigorous graph vector field framework of expressing a vector on a discrete graph, then reconstructing governing full dynamical equations of genome-wide cellular state transitions from snapshot single cell genomics data. This method is complementary to continuous cell state space vector field-based methods (6), and can also be used as a numerical algorithm for solving the high-dimensional continuous equations. Both approaches provide dynamical systems theory-based models to describe the temporal evolution of cell states.

We want to emphasize that both the previous dynamo and current graph-FPE formalisms use estimated RNA velocities, including the splicing-based RNA velocity and various modifications, as well as metabolic labeling data, as inputs. In most splicing-based RNA velocity methods, a velocity is scaled by an unknown splicing rate constant that is in principle different for each gene. Therefore, the magnitude of such RNA velocity vector does not accurately reflect the absolute gene expression rate, and the associated graph FPE model also inherits such lack of quantitative nature. In principle, one may estimate the absolute gene expression with additional information from time course scRNA-seq data. On the other hand, a metabolic labeling-based approach has the capacity of measuring the absolute transcription rate of a gene (up to read count normalization and other data processing procedures (10)), and the velocity has the physical dimension of the number of molecules per unit time. Consequently, the associated graph FPE-based model can provide information on the cell commitment timeline.

Both the dynamo and the graph FPE frameworks assume that the RNA velocity of a cell is related to the transcriptomic state of the cell at the same time, largely restricted by that only snapshot data are used as input. This is a likely strong Markovian assumption, since it takes a finite time for mRNA to be translated into proteins, which are the effectors for gene regulation. Molecular species not explicitly included also contribute to regulations as intermediates, which may also delay regulation. The projection procedure assures the velocity vectors to be in the tangent space of the variable manifold, thus enforces the mathematical self-consistency of the used Markovian dynamics formalism. With information from additional modality especially proteomic data, one can relate the RNA velocity directly to the protein levels. One may also generalize the learning procedure to computationally relax the Markovian approximation by relating the RNA velocity and cell transcriptomic state of different cells.

Most parameters of a graph-FPE model can be determined from the single cell data. While Eqn. 2 provides a procedure for estimating the convection term in Eqn. 8, some alternative and improved methods may be developed in future studies. Different interpretations of the SDEs (Eqn. 5) lead to different forms of the FPE (Eqn. 7) (27), and further studies may examine how well each of these forms models the single cell data. In general, the diffusion constant is gene- and cell state-specific, and the cell birth-death rate is cell state-specific. Determining these parameters requires additional experimental input. One may also examine the chemical Langevin equation formalism (28), in which the convection and the diffusion terms are related. The only adjustable parameter then is the neighborhood size, which should be determined by comparing the relative contributions from convection and diffusion to achieve numerical convergence of the results.

In summary, the present work can be viewed as part of growing efforts spanning various fields of science and engineering of learning dynamical governing equations from high throughput data. As for cell biology, with increasingly sophisticated estimation of RNA velocity (29, 30) and potentially other experimental measurement of gene expression kinetics (31), we expect an increasing amount of quantitative data available that can feed into data-informed dynamical systems theory modeling of the cellular processes. The framework discussed here can be applied to other modality of single cell data, for example, live cell imaging data describing cells in composite cell feature space where velocities can be measured directly (32). It can also be applied to processes other than cell state transitions provided they can be modeled by the FPE formalism.

## Author contributions

YZ and JX conceived and formulated the theoretical framework, YZ performed tests on simulated data, YZ and XQ analyzed scRNA-seq datasets, YZ, XQ, and KN developed the package graph-dynamo, and all authors wrote the manuscript.

## Acknowledgments

We thank Hong Qian and Ping Ao for helpful discussions. This work was partially supported by National Institute of Diabetes and Digestive and Kidney Diseases (R01DK119232), National Institute of General Medical Sciences (1R01GM148525), and National Science Foundation (2325149) to JX, Longevity Impetus Grants to XQ and JSW, and K99 HG012887-01 to XQ.

## Data and Code Availability

Python package graph-dynamo can be accessed from https://github.com/xing-lab-pitt/graph-dynamo.

## References

1. J. Xing, Reconstructing data-driven governing equations for cell phenotypic transitions: integration of data science and systems biology. Physical Biology 19, 061001 (2022).

2. G. H. T. Yeo, S. D. Saksena, D. K. Gifford, Generative modeling of single-cell time series with PRESCIENT enables prediction of cell trajectories with interventions. Nature Communications 12, 3222 (2021).

3. C. Weinreb, S. Wolock, B. K. Tusi, M. Socolovsky, A. M. Klein, Fundamental limits on dynamic inference from single-cell snapshots. Proc Natl Acad Sci U S A 115, E2467–E2476 (2018).

4. G. Schiebinger et al., Optimal-Transport Analysis of Single-Cell Gene Expression Identifies Developmental Trajectories in Reprogramming. Cell 176, 928-943.e922 (2019).

5. G. La Manno et al., RNA velocity of single cells. Nature 10.1038/s41586-018-0414-6 (2018).

6. X. Qiu et al., Mapping Transcriptomic Vector Fields of Single Cells. Cell 185, 690–711 (2022).

7. G. E. Karniadakis et al., Physics-informed machine learning. Nature Reviews Physics 3, 422–440 (2021).

8. S. Chandrasekhar, Stochastic Problems in Physics and Astronomy. Reviews of Modern Physics 15, 1–89 (1943).

9. Z. Chen, W. C. King, A. Hwang, M. Gerstein, J. Zhang, DeepVelo: Single-cell transcriptomic deep velocity field learning with neural ordinary differential equations. Science Advances 8, eabq3745.

10. G. Gorin, M. Fang, T. Chari, L. Pachter, RNA velocity unraveled. PLOS Computational Biology 18, e1010492 (2022).

11. J.-M. Ginoux, Slow Invariant Manifolds of Slow–Fast Dynamical Systems. International Journal of Bifurcation and Chaos 31, 2150112 (2021).

12. L. Cai, N. Friedman, X. S. Xie, Stochastic protein expression in individual cells at the single molecule level. Nature 440, 358–362 (2006).

13. M. Lange et al., CellRank for directed single-cell fate mapping. Nat Methods 19, 159–170 (2022).

14. E. Becht et al., Dimensionality reduction for visualizing single-cell data using UMAP. Nat Biotechnol 1038/nbt.4314 (2018).

15. Y. L. Klimontovich, Nonlinear Brownian motion. Physics-Uspekhi 37, 737 (1994).

16. P. Hänggi, Correlation functions and masterequations of generalized (non-Markovian) Langevin equations. Zeitschrift für Physik B Condensed Matter 31, 407–416 (1978).

17. J. Xing, H.-Y. Wang, G. Oster, From continuum Fokker-Planck models to discrete kinetic models. Biophys. J. 89, 1551–1563 (2005).

18. H. Wang, C. Peskin, T. Elston, A robust numerical algorithm for studying biomolecular transport processes. J. Theo. Biol. 221, 491–511 (2003).

19. L.-H. Lim, Hodge Laplacians on Graphs. SIAM Review 62, 685–715 (2020).

20. X. Qiu, S. Ding, T. Shi, From understanding the development landscape of the canonical fate-switch pair to constructing a dynamic landscape for two-step neural differentiation. PLoS One 7, e49271 (2012).

21. A. Bastidas-Ponce et al., Comprehensive single cell mRNA profiling reveals a detailed roadmap for pancreatic endocrinogenesis. Development 146, dev173849 (2019).

22. R. Yamamoto et al., Clonal Analysis Unveils Self-Renewing Lineage-Restricted Progenitors Generated Directly from Hematopoietic Stem Cells. Cell 154, 1112–1126 (2013).

23. K. Maehara, Y. Ohkawa, Modeling latent flows on single-cell data using the Hodge decomposition. bioRxiv 10.1101/592089, 592089 (2019).

24. C. Voisin, Hodge Theory and Complex Algebraic Geometry I, Cambridge Studies in Advanced Mathematics (Cambridge University Press, Cambridge, 2002), vol. 1.

25. C. Weinreb, S. Wolock, B. K. Tusi, M. Socolovsky, A. M. Klein, Fundamental limits on dynamic inference from single-cell snapshots. Proc Natl Acad Sci U S A (2018).

26. M. Y. Hein, J. S. Weissman, Functional single-cell genomics of human cytomegalovirus infection. Nature Biotechnology 40, 391–401 (2022).

27. J. Xing, Mapping between dissipative and Hamiltonian systems. J. Phys. A: Math. Theor. 43, 375003 (2010).

28. D. T. Gillespie, The chemical Langevin Equation. J. Chem. Phys. 113, 297–306 (2000).

29. Q. Qiu et al., Massively parallel and time-resolved RNA sequencing in single cells with scNT-seq. Nature Methods 17, 991–1001 (2020).

30. N. Battich et al., Sequencing metabolically labeled transcripts in single cells reveals mRNA turnover strategies. Science 367, 1151–1156 (2020).

31. S. G. Rodriques et al., RNA timestamps identify the age of single molecules in RNA sequencing. Nat Biotechnol 39, 320–325 (2021).

32. W. Wang et al., Live-cell imaging and analysis reveal cell phenotypic transition dynamics inherently missing in snapshot data. Science Advances 6, eaba9319 (2020).

