## supplemental text for "Graph-Dynamo: Learning stochastic cellular state transition dynamics from single cell data"

**A. Numerical issues of using Eqn. 1 for tangent space projection**

The relation in Eqn. 1 provides an algorithm for projecting a measured $\mathbf{v}$ onto $T_{\mathbf{x}}\mathcal{M}$ by minimizing $\left\|\mathbf{v}_i - \mathbf{v}_{\parallel}(\mathbf{x}_i)\right\|$, which unfortunately is numerically unstable. The redundancy of the basis vectors leads to coefficients that are not uniquely determined, and failure of the projection. To see the latter, notice that in real data the subspace formed by the displacement vectors only approximates the tangent space $T\mathcal{M}$ locally, and it likely contains small components in the orthogonal space. Then the projection procedure tries to express both $\mathbf{v}_{\parallel}(\mathbf{x}_i)$ and all or part of the remaining $\mathbf{v}_{\perp} = \mathbf{v}_i - \mathbf{v}_{\parallel}(\mathbf{x}_i)$ as a linear combination of $\boldsymbol{\delta}_{ij}$ with some $|\phi_{ij}| \gg 0$ (**Fig. 1B**). We experimented with the idea of removing the redundancy through dimension reduction with PCA, but the numerical results were not satisfactory (**Fig. S1C**), possibly due to the approximation of the curved manifold locally by a Euclidean space with insufficient sampling.

**B. Discrete calculus on graphs**

Before establishing discrete calculus, we first need to define the inner products for scalar functions and vector fields on graphs. Denote a graph as $G(V, E)$, where $V$ is the set of the vertices and $E$

the set of edges. For scalar functions $u$ and $w$ defined on vertices $V$, and vector fields $\boldsymbol{\varphi}$ and $\boldsymbol{\psi}$ defined on edges $E$, the inner products on graphs are defined as (1-3):

$$\langle u, w\rangle_V = \sum_{i\in V} u_i w_i, \tag{S1}$$

$$\langle \boldsymbol{\varphi}, \boldsymbol{\psi}\rangle_E = \frac{1}{2}\sum_{e_{ij}\in E} \varphi_{ij}\psi_{ij}\left|e_{ij}\right|^2, \tag{S2}$$

where the vector field is defined as:

$$\boldsymbol{\varphi}_i = \sum_{j\in\mathcal{N}_i} \varphi_{ij} e_{ij}. \tag{S3}$$

Note that the $1/2$ factor in Eq. S2 prevents double counting when $\varphi_{ij}$ and $\psi_{ij}$ are asymmetrical functions. The additional $\left|e_{ij}\right|^2$ factors in $\langle \boldsymbol{\varphi}, \boldsymbol{\psi}\rangle_E$ is to compensate the fact that $\varphi_{ij}$ and $\psi_{ij}$ does not include the length of the edge. Previous studies have established a series of differential operations for graphs (4), analogous to those on a continuous manifold. For a scalar function $u$ defined on the vertices of $G$, its *derivative* at vertex $i$ along the edge $e_{ij}$ can be defined by treating the function as linear on the edge with respect to a coordinate $y_{ij}$ and parametrizing $e_{ij}$ within an interval $[0, \left|e_{ij}\right|]$ (1-3):

$$u'_{ij} = \frac{\mathrm{d}u}{\mathrm{d}y_{ij}} = \frac{u_j - u_i}{\left|e_{ij}\right|}. \tag{S4}$$

The *gradient* of a scalar function is defined such that the resulting (graph) vector field consists of the derivatives in the direction of each connecting edge $e_{ij}$:

$$\boldsymbol{\nabla} u_i = \sum_{j\sim i} u'\left(e_{ij}\right)\frac{e_{ij}}{\left|e_{ij}\right|} = \sum_{j\sim i}\frac{u_j - u_i}{\left|e_{ij}\right|^2} e_{ij}. \tag{S5}$$

In particular, if a vector field $\boldsymbol{\varphi}_i$is the negative gradient of a scalar function $u$ – which is often called a *potential* in this context – i.e., $-\boldsymbol{\nabla} u_i = \boldsymbol{\varphi}_i$ for all $i \in V$, then the vector field $\boldsymbol{\varphi}$ flows from the sources to the sinks of the potential.

The *divergence* is the adjoint of the gradient, which means that for any scalar function $u$ and vector field $\boldsymbol{\varphi}$, there is:

$$\langle \nabla \cdot \boldsymbol{\varphi}, u \rangle_{\Omega} = -\langle \nabla u, \boldsymbol{\varphi} \rangle_{E}. \tag{S6}$$

Let $u$ be a Kronecker delta function $\delta^k$, such that,

$$\delta_i^k = \begin{cases} 1, & i = k \\ 0, & i \neq k \end{cases} \tag{S7}$$

Then,

$$\langle \nabla \cdot \boldsymbol{\varphi}, \delta^k \rangle_{\Omega} = \sum_{i \in \Omega} (\nabla \cdot \boldsymbol{\varphi})_i \delta_i^k = (\nabla \cdot \boldsymbol{\varphi})_k, \tag{S8}$$

and,

$$-\langle \nabla \delta^k, \boldsymbol{\varphi} \rangle_E = -\frac{1}{2} \sum_{e_{ij} \in E} \frac{\delta_j^k - \delta_i^k}{|e_{ij}|^2} \varphi_{ij} |e_{ij}|^2 = -\frac{1}{2} \left( \sum_{i \in \mathcal{N}_k} \varphi_{ik} + \sum_{j \in \mathcal{N}_k} -\varphi_{kj} \right) = \sum_{i \in \mathcal{N}_k} \frac{\varphi_{ki} - \varphi_{ik}}{2}. \tag{S9}$$

Plugging Eq. S8 and Eq. S9 into Eq. S6 leads to the definition of the graph divergence,

$$(\nabla \cdot \boldsymbol{\varphi})_k = \sum_{i \in \mathcal{N}_k} \frac{\varphi_{ki} - \varphi_{ik}}{2} = \sum_{i \in \mathcal{N}_k} \varphi_{ik}^{\text{asym}}, \tag{S10}$$

where $\varphi_{ij}^{\text{asym}} = (\varphi_{ij} - \varphi_{ji})/2$ denotes the asymmetrical component of the following decomposition of the vector field,

$$\boldsymbol{\varphi}_i = \boldsymbol{\varphi}_i^{\text{sym}} + \boldsymbol{\varphi}_i^{\text{asym}}, \tag{S11}$$

where $\varphi_{ij}^{\text{sym}} = (\varphi_{ij} + \varphi_{ji})/2$. Finally, the divergence of the gradient yields the *graph Laplacian*:

$$\Delta u_i = \nabla \cdot \nabla u_i = \sum_{j \neq i} \frac{1}{|e_{ij}|^2} (u_j - u_i). \tag{S12}$$

Further discussions on the formulation and physical dimensional analysis of graph operators can be found in **Section E**.

#### C. Differential operators for alternative definition of graph vector field

Notice that there is flexibility of choosing the basis vectors $e_{ij}$ while defining a graph vector field. Here we provide an alternative definition of graph vector fields uses normalized edges as bases,

$$\boldsymbol{\varphi}_i = \sum_{j \in \mathcal{N}_i} \varphi_{ij} \frac{e_{ij}}{|e_{ij}|}. \tag{S13}$$

Then the gradient becomes,

$$(\nabla u)_i = \sum_{j\in\mathcal{N}_i} \frac{u_j - u_i}{|e_{ij}|}\frac{e_{ij}}{|e_{ij}|}, \tag{S14}$$

and the inner product on edges is defined as,

$$\langle \boldsymbol{\varphi}, \boldsymbol{\psi} \rangle_E = \frac{1}{2}\sum_{i\in\Omega}\sum_{j\in\mathcal{N}_i} \varphi_{ij}\psi_{ij}. \tag{S15}$$

Using the same approach based on graph Kronecker delta function introduced in **Section B**,ion the divergence becomes,

$$(\nabla \cdot \boldsymbol{\varphi})_i = \sum_{j\in\mathcal{N}_i} \frac{\varphi_{ji} - \varphi_{ij}}{2|e_{ij}|}. \tag{S16}$$

It can be easily verified that the gradient and the divergence derived from the alternative definition of the graph vector field have the same physical dimensions as those from the original definition.

**D. Special considerations for defining dynamics on a graph**

While cell *j* is identified using a specific algorithm to be within the neighborhood of cell *i* and an edge $e_{ij}$exists, the reverse does not necessarily hold using the same algorithm. When a graph is formed on approximating the manifold $\mathcal{M}$, having $e_{ij}$ but not $e_{ji}$ might lead to current formed by diffusion eve without the convection term in Eqn. 11. To avoid such unphysical current, we require that both $e_{ij}$ and $e_{ji}$ exist on the graph when cell *j* is identified as a neighbor of cell *i* and/or cell *i* is identified as a neighbor of cell *i*.

The edge components $\varphi_{ij}$ of a graph vector field has the physical interpretation of convection-contributed net transition probability from vortex *i* to *j*. That is, $\varphi_{ij}$ should be nonnegative. In practice, the relation might not be satisfied since the two terms are obtained from $v_i$ and $v_j$, respectively. Therefore, we imposed this constraint on the drift term used in the graph FPE as follows,

$$\varphi_{ij}^{\mathrm{FPE}} = \begin{cases} \|\varphi_{ij} - \varphi_{ji}\|/2, & \varphi_{ij} \geq \varphi_{ji} \\ 0, & \varphi_{ij} < \varphi_{ji} \end{cases} \tag{S17}$$

For simplicity, we omit the superscript for the drift term in graph FPE, but it should be understood as the net velocity flow calculated from the graph vector field.

**E. The chemical master equation, Gillespie algorithm, and graph FPE**

Consider a system of $N$ species, whose state is defined as the number of molecules for all species, $\mathbf{x} = [x_1, x_2, \dots, x_N]$, and $M$ reactions. The system state changes through chemical reactions prescribed by the *stoichiometry matrix* $[\boldsymbol{\delta}_1, \boldsymbol{\delta}_2, \dots, \boldsymbol{\delta}_M]$, with each $\boldsymbol{\delta}_m$ encodes how the number of molecules for each species change after reaction $m$. Then a set of the chemical master equations (CMEs) depict how the probability of the state of the system evolves (3, 5):

$$\frac{\partial p(\mathbf{x}, t)}{\partial t} = \sum_{m=1}^{M} a_m(\mathbf{x} - \boldsymbol{\delta}_m) p(\mathbf{x} - \boldsymbol{\delta}_m, t) - a_m(\mathbf{x}) p(\mathbf{x}, t) \qquad \text{(S18)}$$

The function $a(\mathbf{x})$, called the *propensity*, gives the probability of occurrence of reaction $j$ per unit time. Given a stoichiometry matrix, propensity functions, and initial conditions, one may use the Gillespie algorithm to generate trajectories that are exact samples of the solution of the corresponding CME (5).

By comparing Eq. S18 and Eq. 8 in the main text, the graph FPE has the same form as the CME for cell $i$, whose edges connected to $k$ neighbors are the "reactions", and the corresponding distance vectors $[\boldsymbol{\delta}_{i1}, \boldsymbol{\delta}_{i2}, \dots, \boldsymbol{\delta}_{ik}]$ form the "stoichiometry matrix", with an additional diffusion term. Although often under various steps of preprocessing, an RNA velocity vector can be regarded as a vector of reaction rates, defined as the number of molecules generated per unit time. It is connected to the propensity by:

**Deleted:** is locally

**Deleted:** a

**Deleted:** the

$$\mathbf{v} = \lim_{\Delta t \to 0} \frac{\mathbb{E}[\Delta \mathbf{x} | \mathbf{x}]}{\Delta t} = \lim_{\Delta t \to 0} \frac{1}{\Delta t} \sum_{m=1}^{M} \boldsymbol{\delta}_m \operatorname{Pr}_m(\mathbf{x}) = \sum_{m=1}^{M} \boldsymbol{\delta}_m \, a_m(\mathbf{x}) \qquad \text{(S19)}$$

where $\operatorname{Pr}_m(\mathbf{x})$ is the probability for the reaction $m$ to occur within an infinitesimal time interval $\Delta t$, and by definition $\operatorname{Pr}_m(\mathbf{x}) = \lim_{\Delta t \to 0} a_m(\mathbf{x}) \Delta t$. Comparing Eq. S19 and Eq. S3, the coefficient of the graph vector field $\varphi_{ij}$ models the propensity for the transition from state $\mathbf{x}_i$ to $\mathbf{x}_j$, through the "stoichiometry vector" $\boldsymbol{\delta}_{ij}$. Eq. S19 also provides an alternative rationale for representing RNA velocity vector using the graph vector field from the perspective of CME. Compared to the geometrical explanation, Eq. S19 has the advantage of generalization in both continuous and discrete domains, and gene expression models are intrinsically discrete. Therefore, it is even desirable to estimate RNA velocity directly using algorithms based on Eq. S19, as previously discussed (6). The diffusion term in the graph FPE represents various sources of noises in addition to the intrinsic stochasticity of gene expressions, which is now a tuning parameter for the width of the probability distribution from the solution of Eq. 9.

The pipeline of setting up Gillespie simulations and conforming simulated data format to the AnnData format (7) is implemented as a simulation module in ***dynamo***.

**F. ODEs and the Gillespie simulation of the two-gene toggle switch model**

We used activation and inhibition Hill functions to model the induction and suppression effects between the two genes (see **Fig. 2A** in the main text for the diagram of the model):

$$x_1 = \frac{a_1 x_1^n}{S_1^n + x_1^n} + \frac{b_1 K_1^n}{K_1^n + x_2^n} - \gamma_1 x_1, \quad \text{(S20)}$$

$$x_2 = \frac{a_2 x_2^n}{S_2^n + x_2^n} + \frac{b_2 K_2^n}{K_2^n + x_1^n} - \gamma_2 x_2. \quad \text{(S21)}$$

To control the size (number of molecules) and timescale of the simulation, we introduced two additional parameters, $r$ and $\tau$, such that for $i \in \{1,2\}$, $a_i = \tilde{a}_i r/\tau$, $b_i = \tilde{b}_i r/\tau$, $S_i = \tilde{S}_i r$, $K_i = \tilde{K}_i r$, and $\gamma_i = \tilde{\gamma}_i/\tau$. The fixed points of this system can only be solved numerically, and depending on the parameters, there could be 1, 2, and 3 stable fixed points (attractors) in total (see Figure xxx for the bifurcation plot). To generate data with a clear bifurcation of two attractors, we used the following parameters: $a_1 = a_2 = b_1 = b_2 = 0.5$, $\gamma_1 = \gamma_2 = 0.2$, $S_1 = S_2 = K_1 = K_2 = 2.5$, $n = 5$. In this parameter regime, the two attractors are symmetrically distributed near the *x*- and *y*-axis, respectively. For the attractor near the *x*-axis, $x_2 \ll K_1$, $x_1 \gg S_1$, and because at the fixed point the velocities are zero, i.e., $x_1 = 0$, from Eq. S20:

$$\begin{aligned} &0 \sim a_1 + b_1 - \gamma_1 x_1^{\mathrm{ss}} \\ &\rightarrow x_1^{\mathrm{ss}} \sim \frac{a_1 + b_1}{\gamma_1}. \end{aligned} \quad \text{(S22)}$$

where $x_1^{\mathrm{ss}}$ stands for the steady-state value of $x_1$ at the attractor near the *x*-axis. By symmetry, we also have $x_2^{\mathrm{ss}} \sim (a_2 + b_2)/\gamma_2$ for the attractor near the *y*-axis. We placed ten cells at $(x_1^{\mathrm{ss}}, x_2^{\mathrm{ss}})$, each added with a Gaussian noise ($\sigma = 5$), as the initial conditions for the ten simulated trajectories.

For each gene, we divide the ODE into propensities for two reactions: synthesis and degradation. For gene $i$, the stoichiometry for the degradation is -1, i.e., whenever a degradation reaction happens, the number of molecules for gene $i$ is reduced by one. Similarly, the stoichiometry for the synthesis is +1 for gene $i$. The propensity for the degradation, based on Eq. S19, is:

$$a_{\text{deg}}(x_i) = \frac{v_{\text{deg}}(x_i)}{\delta_{\text{deg}}^i} = -\gamma_i x_i, \tag{S23}$$

where the subscript "deg" stands for "degradation", $v$ is the reaction rate (from the ODE), and $\delta_{\text{deg}}^i$ the stoichiometry of gene $i$ for the degradation. For the synthesis, the propensity is the sum of self-activation and mutual inhibition,

$$a_{\text{syn}}(x_i) = \frac{v_{\text{syn}}(x_i)}{\delta_{\text{syn}}^i} = \frac{a_i x_i^n}{S_i^n + x_i^n} + \frac{b_i K_i^n}{K_i^n + x_j^n}, \tag{S24}$$

where $j$ is the other gene than gene $i$.

**G. ODEs and the Gillespie simulation of the neurogenesis model**

Based on the work of Qiu et al. (8), the ODEs for the 12 genes involved in the neurogenesis model are defined as:

$$x_{\text{pax6}} = \frac{a_{\text{pax6}} K_{\text{pax6}}^n}{K_{\text{pax6}}^n + x_{\text{tuj1}}^n + x_{\text{a1dh1l}}^n + x_{\text{sox8}}^n} - \gamma_{\text{pax6}} x_{\text{pax6}}, \tag{S25}$$

$$x_{\text{mash1}} = \frac{a_{\text{mash1}} x_{\text{pax6}}^n}{K_{\text{mash1}}^n + x_{\text{pax6}}^n + x_{\text{hes5}}^n} - \gamma_{\text{mash1}} x_{\text{mash1}}, \tag{S26}$$

$$x_{\text{zic1}} = \frac{a_{\text{zic1}} x_{\text{mash1}}^n}{K_{\text{zic1}}^n + x_{\text{mash1}}^n} - \gamma_{\text{zic1}} x_{\text{zic1}}, \tag{S27}$$

$$x_{\text{brn2}} = \frac{a_{\text{brn2}} x_{\text{mash1}}^n}{K_{\text{brn2}}^n + x_{\text{mash1}}^n + x_{\text{olig2}}^n} - \gamma_{\text{brn2}} x_{\text{brn2}}, \tag{S28}$$

$$x_{\text{tuj1}} = \frac{a_{\text{tuj1}} \left(x_{\text{zic1}}^n + x_{\text{brn2}}^n + x_{\text{myt1l}}^n\right)}{K_{\text{tuj1}}^n + x_{\text{zic1}}^n + x_{\text{brn2}}^n + x_{\text{myt1l}}^n} - \gamma_{\text{tuj1}} x_{\text{tuj1}}, \tag{S29}$$

$$x_{\text{hes5}} = \frac{a_{\text{hes5}} x_{\text{pax6}}^n}{K_{\text{hes5}}^n + x_{\text{pax6}}^n + x_{\text{mash1}}^n} - \gamma_{\text{hes5}} x_{\text{hes5}}, \tag{S30}$$

$$x_{\text{scl}} = \frac{a_{\text{scl}} x_{\text{hes5}}^n}{K_{\text{scl}}^n + x_{\text{hes5}}^n + x_{\text{olig2}}^n} - \gamma_{\text{scl}} x_{\text{scl}}, \tag{S31}$$

$$x_{\text{olig2}} = \frac{a_{\text{olig2}} x_{\text{hes5}}^n}{K_{\text{olig2}}^n + x_{\text{hes5}}^n + x_{\text{scl}}^n} - \gamma_{\text{olig2}} x_{\text{olig2}}, \tag{S32}$$

$$x_{\mathrm{stat3}} = \frac{a_{\mathrm{stat3}} x_{\mathrm{hes5}}^n x_{\mathrm{scl}}^n}{K_{\mathrm{stat3}}^n + x_{\mathrm{hes5}}^n x_{\mathrm{scl}}^n} - \gamma_{\mathrm{stat3}} x_{\mathrm{stat3}}, \tag{S33}$$

$$x_{\mathrm{a1dh1l}} = \frac{a_{\mathrm{a1dh1l}} x_{\mathrm{stat3}}^n}{K_{\mathrm{a1dh1l}}^n + x_{\mathrm{stat3}}^n} - \gamma_{\mathrm{a1dh1l}} x_{\mathrm{a1dh1l}}, \tag{S34}$$

$$x_{\mathrm{myt1l}} = \frac{a_{\mathrm{myt1l}} x_{\mathrm{olig2}}^n}{K_{\mathrm{myt1l}}^n + x_{\mathrm{olig2}}^n} - \gamma_{\mathrm{myt1l}} x_{\mathrm{myt1l}}, \tag{S35}$$

$$x_{\mathrm{sox8}} = \frac{a_{\mathrm{sox8}} x_{\mathrm{olig2}}^n}{K_{\mathrm{sox8}}^n + x_{\mathrm{olig2}}^n} - \gamma_{\mathrm{sox8}} x_{\mathrm{sox8}}. \tag{S36}$$

Intuitively, the above ODEs contain two consecutive toggle switch motifs (see **Fig. 2B** in the main text for the diagram): the mutual inhibition between *hes5* and *mash1*, both activated by *pax6*, and the mutual inhibition between *scl* and *olig2*, both activated by *hes5*. The first toggle switch creates the branch for the neuron (high *mash1* and low *hes5* expression), and the second toggle switch creates the branch for the astrocyte (high *scl*) and the oligodendrocyte (high *olig2*). The purpose of this simulation is not to mimic exactly the biological neurogenesis, but to provide three stable cell types with different commitment timelines to demonstrate the effectiveness of our method. Like the two-gene toggle switch model, we introduced the size and timescale parameter $r$ and $\tau$ for fine adjustments. The strategy for creating the stoichiometry matrix and the propensity function is the same as the two-gene toggle switch model, i.e., for each gene the reaction is divided into the synthesis and degradation with simple stoichiometry, and the propensity for each is calculated based on Eq. S23 and Eq. S24. We used the following parameter sets to create the three attractors: $a_{\mathrm{pax6}} = 2.2$, $a_{\mathrm{mash1}} = a_{\mathrm{hes5}} = 4$, $a_{\mathrm{zic1}} = a_{\mathrm{brn2}} = a_{\mathrm{tuj1}} = a_{\mathrm{stat3}} = a_{\mathrm{a1dh1l}} = a_{\mathrm{myt1l}} = a_{\mathrm{sox8}} = 3$, $a_{\mathrm{scl}} = a_{olig2} = 5$. $K = 1$ for all genes, except that $K_{\mathrm{pax6}} = 1$. $\gamma = 1$, $n = 4$ for all genes. All genes have zero number of molecules at the beginning of the simulation.

#### H. Derivation of Mean Exit Time and Mean First Passage Time for Simulated Trajectories and Continuous Time Markov Chains

The sink states of a Markov chain are those whose outward transition rates are zero, so that any cell that reaches the sink states is trapped. Denote the set containing all sink states as $\mathcal{S}$, the Mean Exit Time (MET) is defined as the expected time for the cell to reach *any* sink state from the initial position for the first time. For simulated trajectories, the MET can be computed based on the definition directly:

$$\langle \tau_{\text{exit}} \rangle = \frac{1}{n_{\mathcal{S}}} \sum_{i=1}^{n_{\mathcal{S}}} (t_{\text{sink}} - t_{\text{init}}) \tag{S37}$$

where $t_{\text{sink}}$ is the simulation time for the trajectory to reach the sink set $\mathcal{S}$ immediately after the trajectory reaches the initial state at $t_{\text{init}}$. In our simulations, the initial state is defined as the collection of cells whose simulation time $t < 1$. $\mathcal{S}$ contains cells in circles of the two attractors with a radius of 5 for the two-gene model, and three attractors with a radius of 10 for the neurogenesis model. $n_{\mathcal{S}}$ is the number of trajectories that have reached any state in $\mathcal{S}$. If we are interested in the time for cells to reach a subset of $\mathcal{S}$, denoted as the target set $\mathcal{T}$, we can define the Mean First Passage Time (MFPT) as the expected time for a cell to reach $\mathcal{T}$, before getting trapped by any other sinks in $\mathcal{S}$. Similar to the MET, the MFPT of simulated trajectories can be calculated directly based on the definition:

$$\langle \tau_{\text{pass}} \rangle = \frac{1}{n_{\mathcal{T}\backslash\mathcal{S}}} \sum_{i=1}^{n_{\mathcal{T}\backslash\mathcal{S}}} \left(t_{\text{target}} - t_{\text{init}}\right) \tag{S38}$$

where $n_{\mathcal{T}\backslash\mathcal{S}}$ is the number of trajectories that have reached any state in $\mathcal{T}$ without first reaching other states in $\mathcal{S}$.

For the CTMC constructed based on the graph FPE, given the transition rate matrix $\mathbf{Q}$, initial distribution vector $\mathbf{p}_0$, and the sink states $\mathcal{S}$, the MET can be calculated by (9):

$$\langle \tau_{\text{exit}} \rangle = -\sum_{i} \left[\mathbf{Q}^{-1}\mathbf{p}_0\right]_i, \tag{S39}$$

where $\mathbf{Q}$ is a submatrix of $\mathbf{Q}$, whose columns and rows of the sink states are removed, and similarly, $\mathbf{p}_0$ is the initial distribution vector with removed target state elements. The MFPT for CTMCs, given the target states $\mathcal{T}$ and other sink states $\mathcal{S}$, can be calculated as (9, 10):

$$\langle \tau_{\text{pass}} \rangle = -\frac{\sum_{j\in\mathcal{T}} \sum_{i\notin\mathcal{S}} q_{i\to j} \left[\mathbf{Q}^{-2}\mathbf{p}_0\right]_i}{\sum_{j\in\mathcal{T}} \sum_{i\notin\mathcal{S}} q_{i\to j} \left[\mathbf{Q}^{-1}\mathbf{p}_0\right]_i} \tag{S40}$$

where $\mathbf{Q}$ is the submatrix of $\mathbf{Q}$ with removed columns and rows of states in the set $\mathcal{T}$ or $\mathcal{S}$, and $\mathbf{p}_0$ the vector with removed elements of target and exiting states.

To see how Eq. S39 and Eq. S40 can be derived from the definitions of the MET and MFPT, because sinks absorb any trajectory, in the sense that the transition rate, denoted as $q_{i\to j}$, from the sink to any other vertex is 0, i.e., for any $i \in \mathcal{S}$ and $j \notin \mathcal{S}$:

$$q_{i\to j} = Q_{ji} = 0 \tag{S41}$$

The probability for a trajectory to not enter any sink vertex before $t$ and transition into the target set $\mathcal{T}$ in the infinitesimal time interval $[t, t + \mathrm{d}t]$ is the following quantity up to a normalization factor:

$$\sum_{i\notin\mathcal{S}} \sum_{j\in\mathcal{T}} p_i(t) q_{i\to j} \mathrm{d}t, \tag{S42}$$

where $p_i(t)$ is the probability that the trajectory is at vertex $i$ at $t$, satisfying the master equation (Eq. S18). The normalization factor is:

$$\begin{aligned} Z &= \int_0^\infty \sum_{i\notin\mathcal{S}} \sum_{j\in\mathcal{T}} p_i(t) q_{i\to j} \mathrm{d}t \\ &= \int_0^\infty \sum_{i\notin\mathcal{S}} \left[e^{\mathbf{Q}t}\mathbf{p}_0\right]_i \sum_{j\in\mathcal{T}} q_{i\to j} \mathrm{d}t \\ &= \sum_{i\notin\mathcal{S}} \left[\left(e^{\mathbf{Q}t}\Big|_0^\infty\right) \mathbf{Q}^{-1}\mathbf{p}_0\right]_i \sum_{j\in\mathcal{T}} q_{i\to j} \\ &= -\sum_{j\in\mathcal{T}} \sum_{i\notin\mathcal{S}} q_{i\to j} \left[\mathbf{Q}^{-1}\mathbf{p}_0\right]_i. \end{aligned} \tag{S43}$$

Given the above probability distribution, the MFPT is the following expectation:

$$\begin{aligned} \langle \tau_{\text{pass}} \rangle &= \frac{1}{Z} \int_0^\infty t \sum_{i\notin\mathcal{S}} \sum_{j\in\mathcal{T}} p_i(t) q_{i\to j} \mathrm{d}t \\ &= \frac{1}{Z} \int_0^\infty t \sum_{i\notin\mathcal{S}} \sum_{j\in\mathcal{T}} \left[e^{\mathbf{Q}t}\mathbf{p}_0\right]_i q_{i\to j} \mathrm{d}t \\ &= \frac{1}{Z} \sum_{i\notin\mathcal{S}} \sum_{j\in\mathcal{T}} q_{i\to j} \left[\left( t e^{\mathbf{Q}t}\Big|_0^\infty - \int_0^\infty e^{\mathbf{Q}t}\mathrm{d}t \right) \mathbf{Q}^{-1}\mathbf{p}_0\right]_i \end{aligned}$$

$$= \frac{1}{Z}\sum_{i\notin\mathcal{S}}\sum_{j\in\mathcal{T}} q_{i\to j}\left[\left(-e^{\mathbf{Q}t}\Big|_0^\infty\right)\mathbf{Q}^{-2}\mathbf{p}_0\right]_i$$

$$= -\frac{\sum_{j\in\mathcal{T}}\sum_{i\notin\mathcal{S}} q_{i\to j}\left[\mathbf{Q}^{-2}\mathbf{p}_0\right]_i}{\sum_{j\in\mathcal{T}}\sum_{i\notin\mathcal{S}} q_{i\to j}\left[\mathbf{Q}^{-1}\mathbf{p}_0\right]_i}. \qquad \text{(S44)}$$

When $\mathcal{T} = \mathcal{S}$, the target set is the only exit for the Markov chain, and the above equation can be reduced to that for the MET. In this case, the normalization factor sums to one:

$$Z = \int_0^\infty \sum_{i\notin\mathcal{S}}\sum_{j\in\mathcal{T}} p_i(t) q_{i\to j}\mathrm{d}t = \int_0^\infty \sum_{j\in\mathcal{T}} \frac{\mathrm{d}p_j(t)}{\mathrm{d}t}\mathrm{d}t = \sum_{j\in\mathcal{T}} p_j(t)\Big|_0^\infty = 1. \qquad \text{(S45)}$$

In the first step, we used the fact that, for sink states $j \in \mathcal{T}$, there is no transition rate out from the state, so from the master equation:

$$\frac{\mathrm{d}p_j(t)}{\mathrm{d}t} = \sum_{i\notin\mathcal{S}} p_i(t) q_{i\to j}, \qquad \text{(S46)}$$

and in the final step, we used the fact that for $j \in \mathcal{T}$, $p_j(0) = 0$, and $\sum_{j\in\mathcal{T}} p_j(\infty) = 1$ because all trajectories will be absorbed into the sink states eventually. Therefore, the expression for MET becomes:

$$\langle \tau_{\mathrm{exit}} \rangle = \int_0^\infty t \sum_{i\notin\mathcal{S}}\sum_{j\in\mathcal{T}} p_i(t) q_{i\to j}\mathrm{d}t$$

$$= \int_0^\infty t \sum_{j\in\mathcal{T}} \frac{\mathrm{d}p_j(t)}{\mathrm{d}t}\mathrm{d}t$$

$$= \sum_{j\in\mathcal{T}}\left(p_j(t)t\Big|_0^\infty - \int_0^\infty p_j(t)\mathrm{d}t\right)$$

$$= -\sum_{j\in\mathcal{T}}\int_0^\infty \left[e^{\mathbf{Q}t}\mathbf{p}_0\right]_j \mathrm{d}t$$

$$= -\sum_{j\in\mathcal{T}}\left[\mathbf{Q}^{-1}\mathbf{p}_0\right]_j. \qquad \text{(S47)}$$

**I. State Lumping for Discrete and Continuous Time Markov Chains**

Let $\mathcal{A}$ and $\mathcal{B}$ be two subsets of vertices in the transition graph $G$. For discrete time Markov chains, the transition probability from some vertex $i \in \mathcal{A}$ to all vertices in $\mathcal{B}$ is:

$$p_{i\to\mathcal{B}} = P(\{j|j \in \mathcal{B}\}|i) = \frac{\sum_{j\in\mathcal{B}} P(j|i)p_i(t)}{p_i(t)} = \sum_{j\in\mathcal{B}} P(j|i) = \sum_{j\in\mathcal{B}} p_{i\to j}, \quad \text{(S48)}$$

which is independent of the state probability distribution $p_i(t)$. However, the overall transition probability from all vertices in $\mathcal{A}$ to those in $\mathcal{B}$ is a function of state probabilities $p_i(t)$ for vertices in $\mathcal{A}$:

$$p_{\mathcal{A}\to\mathcal{B}} = P(\{j|j \in \mathcal{B}\}|\{i|i \in \mathcal{A}\}) = \frac{\sum_{i\in\mathcal{A}}\sum_{j\in\mathcal{B}} p_{i\to j}p_i(t)}{\sum_{i\in\mathcal{A}} p_i(t)} = \frac{\sum_{i\in\mathcal{A}} p_{i\to\mathcal{B}}p_i(t)}{\sum_{i\in\mathcal{A}} p_i(t)}. \quad \text{(S49)}$$

Therefore, the transition dynamics of the lumped Markov chain is no longer Markovian, and the transition probability $p_{\mathcal{A}\to\mathcal{B}}$ changes as the state probability propagates over time. However, when the original Markov chain is at steady state, i.e., $p_i(t) = \pi_i$ for $i \in \mathcal{A}$, the transition probability becomes constant:

$$p_{\mathcal{A}\to\mathcal{B}} = \frac{\sum_{i\in\mathcal{A}} p_{i\to\mathcal{B}}\pi_i}{\sum_{i\in\mathcal{A}} \pi_i} \quad \text{(S50)}$$

Then we can obtain the lumped transition probability matrix **P** based on any partition of the vertices. For the transition rate matrix of a continuous time Markov chain, similar to Eq. S50, the lumped transition rate is defined as:

$$q_{\mathcal{A}\to\mathcal{B}} = \frac{\sum_{i\in\mathcal{A}} q_{i\to\mathcal{B}}\pi_i}{\sum_{i\in\mathcal{A}} \pi_i} \quad \text{(S51)}$$

### Supplementary Figures

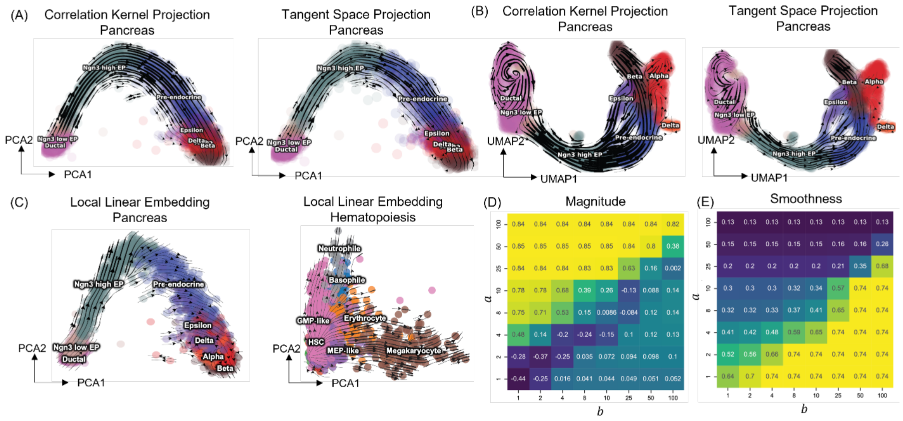


**Figure S1. Tangent space projection (TSP) kernel projection and graph based stochastic dynamics modeling of the hematopoiesis and pancreatic endogenesis dataset.** (A, B) RNA velocity projected to PCA (A) and UMAP (B) using correlation kernel (*left*) and TSP kernel (*right*), visualized as streamline plots for the pancreatic endogenesis dataset. (C) RNA velocity projected using local linear embedding by sampling neighborhoods of 500 datapoints, visualized as streamline plots for the pancreatic endogenesis (*left*) and hematopoiesis dataset (*right*). (D) Heatmap of the correlation of the magnitude with respect to unprojected RNA velocity (*left*), and local smoothness (*right*) of TSP kernel-based RNA velocity projection to UMAP (left) or PCA (right) space under different parameters of $b$ in x-axis and $a$ for the pancreatic endogenesis dataset.

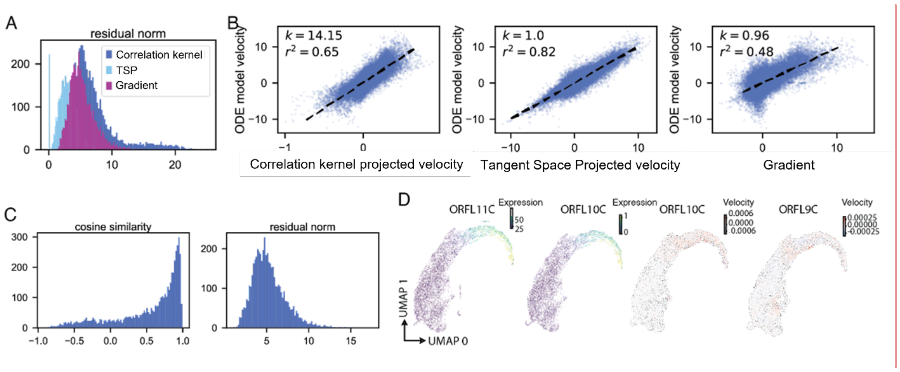


**Fig. S2. Benchmark and application of the graph vector field based gradient operator approach on the simulated neurogenesis and experimental CMV infection dataset.** (A) The distribution of the norm of the residual of the projected RNA velocity in the gene space from three different kernels to the ground truth velocity vectors. (B) Scatterplots of the correlation kernel based (left), TSP kernel based (middle), and gradient based RNA velocities (right) and the true RNA velocities (y-axis) projected onto the original gene expression dimensions across cells. (C) The distribution of cosine similarity and residual norm between the graph operator projected RNA velocity to the ground truth velocity in the original gene expression space. (D) The scatter plot of the gene expression and predicted RNA velocity across cells with the graph gradient operator of viral genes ORFL11C, ORFL10C and ORFL9C.

1. R. W. Feres, Matt, Reaction-diffusion on metric graphs and conversion probability. *arXiv* arXiv:1501.06976 (2015).
2. M. I. Freidlin, A. D. Wentzell, Diffusion Processes on Graphs and the Averaging Principle. *The Annals of Probability* **21**, 2215-2245, 2231 (1993).
3. S.-i. Goto, H. Hino, Diffusion equations from master equations—A discrete geometric approach. *Journal of Mathematical Physics* **61**, 113301 (2020).
4. L.-H. Lim, Hodge Laplacians on Graphs. *SIAM Review* **62**, 685-715 (2020).
5. D. T. Gillespie, Stochastic simulation of chemical kinetics. *Annu Rev Phys Chem* **58**, 35-55 (2007).
6. G. Gorin, M. Fang, T. Chari, L. Pachter, RNA velocity unraveled. *bioRxiv* 10.1101/2022.02.12.480214, 2022.2002.2012.480214 (2022).
7. D. Bredikhin, I. Kats, O. Stegle, MUON: multimodal omics analysis framework. *Genome Biology* **23**, 42 (2022).
8. X. Qiu, S. Ding, T. Shi, From Understanding the Development Landscape of the Canonical Fate-Switch Pair to Constructing a Dynamic Landscape for Two-Step Neural Differentiation. *PLoS ONE* **7**, e49271 (2012).

**Commented [JX1]:** Change “co-optimization” to “Tangent space projection”

9. N. F. Polizzi, M. J. Therien, D. N. Beratan, Mean First-Passage Times in Biology. *Israel Journal of Chemistry* **56**, 816-824 (2016).
10. I. V. Gopich, A. Szabo, Theory of the statistics of kinetic transitions with application to single-molecule enzyme catalysis. *J. Chem. Phys.* **124**, 154712 (2006).
